# A human *in vitro* neuronal model for studying homeostatic plasticity at the network level

**DOI:** 10.1101/2023.04.14.536851

**Authors:** Xiuming Yuan, Sofía Puvogel, Jon-Ruben van Rhijn, Anna Esteve-Codina, Mandy Meijer, Simon Rouschop, Eline J.H. van Hugte, Astrid Oudakker, Chantal Schoenmaker, Dirk Schubert, Barbara Franke, Nael Nadif Kasri

## Abstract

Mechanisms that underlie homeostatic plasticity have been extensively investigated at single-cell levels in animal models, but are less well understood at the network level. Here, we used microelectrode arrays to characterize neuronal networks following induction of homeostatic plasticity in human induced pluripotent stem cell (hiPSC)-derived glutamatergic neurons co-cultured with rat astrocytes. Chronic suppression of neuronal activity through tetrodotoxin (TTX) elicited a time-dependent network re-arrangement. Increased expression of AMPA receptors and the elongation of axon initial segments were associated with increased network excitability following TTX treatment. Transcriptomic profiling of TTX-treated neurons revealed up-regulated genes related to extracellular matrix organization, while down-regulated genes related to cell communication; also astrocytic gene expression was found altered. Overall, our study shows that hiPSC-derived neuronal networks provide a reliable *in vitro* platform to measure and characterize homeostatic plasticity at network and single-cell level; this platform can be extended to investigate altered homeostatic plasticity in brain disorders.

## Introduction

In the healthy brain, neuronal activity adapts dynamically to a changing environment. Two principal mechanisms accommodate such adaptive behaviour: Hebbian plasticity and homeostatic plasticity. Traditional forms of Hebbian plasticity, such as long-term potentiation (LTP) and long-term depression (LTD), induce changes in the strength of individual synaptic connections and constitute the biological substrate of learning and memory consolidation. However, without effective negative feedback regulation, their effects could cause a destabilization of neuronal networks (Fox and Stryker, 2017). Homeostatic plasticity is a negative feedback mechanism that stabilizes neuronal network activity by adjusting synaptic strength and intrinsic properties of the neurons in response to activity perturbations during learning and development. Eventually, the homeostatic changes that occur through these mechanisms prevent the neuronal networks from becoming hypo- or hyper-active (Tien and Kerschensteiner, 2018; Wefelmeyer et al., 2016; Turrigiano, 2012; Watt and Desai, 2010). As such, homeostatic plasticity mechanisms play a critical role in the correct functioning of the nervous system (Antoine et al., 2019). Previous studies identified disrupted homeostatic plasticity in rodent models for different neurodevelopmental disorders (NDDs), such as Fragile X syndrome, Kleefstra syndrome, Rett syndrome, and tuberous sclerosis, suggesting that altered or insufficient homeostatic plasticity during development contributes to cognitive and behavioural impairments that characterize NDDs (Bulow et al., 2019; Lee et al., 2018; Benevento et al., 2016; Dani et al., 2005; Zeng et al., 2007: Antoine et al., 2019).

While homeostatic plasticity mechanisms have been well characterized at the single-cell level, using rodent dissociated-cell cultures and organotypic slice cultures (Moulin et al., 2020; Debanne et al., 2019; Debanne and Russier, 2019), homeostatic plasticity at network level is poorly understood. In addition, it remains unclear how intrinsic properties and synaptic strength cooperate to stabilize neuronal networks in response to changes in activity levels (Antoine et al., 2019; Turrigiano, 2011). Thus, establishing a framework to assess homeostatic plasticity at network level in a human model is essential for a better understanding of human neurodevelopment and neuronal function, both in normal and pathological conditions.

Human induced pluripotent stem cell (hiPSC)-derived neuronal models allow for the generation of networks of controllable cellular composition that contain spontaneously active and synaptically connected neurons in a patient-specific background (Frega et al., 2017; Bardy et al., 2016). They are increasingly used as a model system to understand the pathophysiology of brain disorders, primarily by studying spontaneous activity under basal conditions (McCready et al., 2022; van Hugte and Nadif Kasri, 2019). However, despite the overwhelming implication of synaptic and homeostatic plasticity deficits in brain disorders, only a handful of studies have investigated mechanisms of plasticity in hiPSC-derived neurons (Cordella et al., 2022; Pre et al., 2022; Meijer et al., 2019; Zhang et al., 2018). In particular, no attention has been given to the study of homeostatic plasticity at the network level in hiPSC-derived neurons.

Here, we established a model of tetrodotoxin (TTX)-induced homeostatic plasticity in co-cultures of hiPSC-derived glutamatergic neurons on microelectrode arrays (MEAs), which we have previously shown to facilitate non-invasive, reproducible, real-time, and multidimensional measurement of activity in hiPSC-derived neuronal networks (Mossink et al., 2021). We characterized single-cell and network changes induced with this form of homeostatic plasticity, together with changes in gene expression that may underlie them. We show that hiPSC-derived neuronal networks form a reliable platform to measure and characterize homeostatic plasticity, which can also be harnessed to investigate homeostatic plasticity in human models for brain disorders at the network and single-cell level.

## Results

### TTX-induced homeostatic plasticity leads to re-arrangement of neuronal networks

Populations of human induced pluripotent stem cell (hiPSC)-derived neurons cultured on MEAs generate highly synchronous network bursting activity within a few days *in vitro* (DIV; Frega et al., 2019; Frega et al., 2017). To investigate the effect of chronic activity perturbation on neuronal network dynamics, we co-cultured glutamatergic neurons, derived from three independent control hiPSC lines (Ctrl1, 2, and 3), with rat astrocytes on MEAs and treated them with tetrodotoxin (TTX), a sodium channel blocker, or with a control vehicle (Figure 1A). This procedure allows testing if neuronal networks adapt to changes in neuronal activity by means of homeostatic plasticity. Already at DIV 30, the neuronal networks recorded on MEAs typically exhibited three distinctive patterns of activity: (a) random spiking activity (i.e., isolated a-synchronous action potentials), (b) activity that was self-organized into a local burst (i.e., high frequency trains of spikes), and (c) network bursting activity (i.e., bursts detected in most of the channels; Figure 1B and S1A). At both DIV 30 and DIV 49, the presence of rhythmic and synchronous network bursts, integrated by many spikes and involving most of the channels (Figure 1C, S1B, and S1E), indicated that control hiPSC-derived glutamatergic neurons had organized into synaptically connected and spontaneously active neuronal networks (Frega et al., 2017).

**Figure 1.**
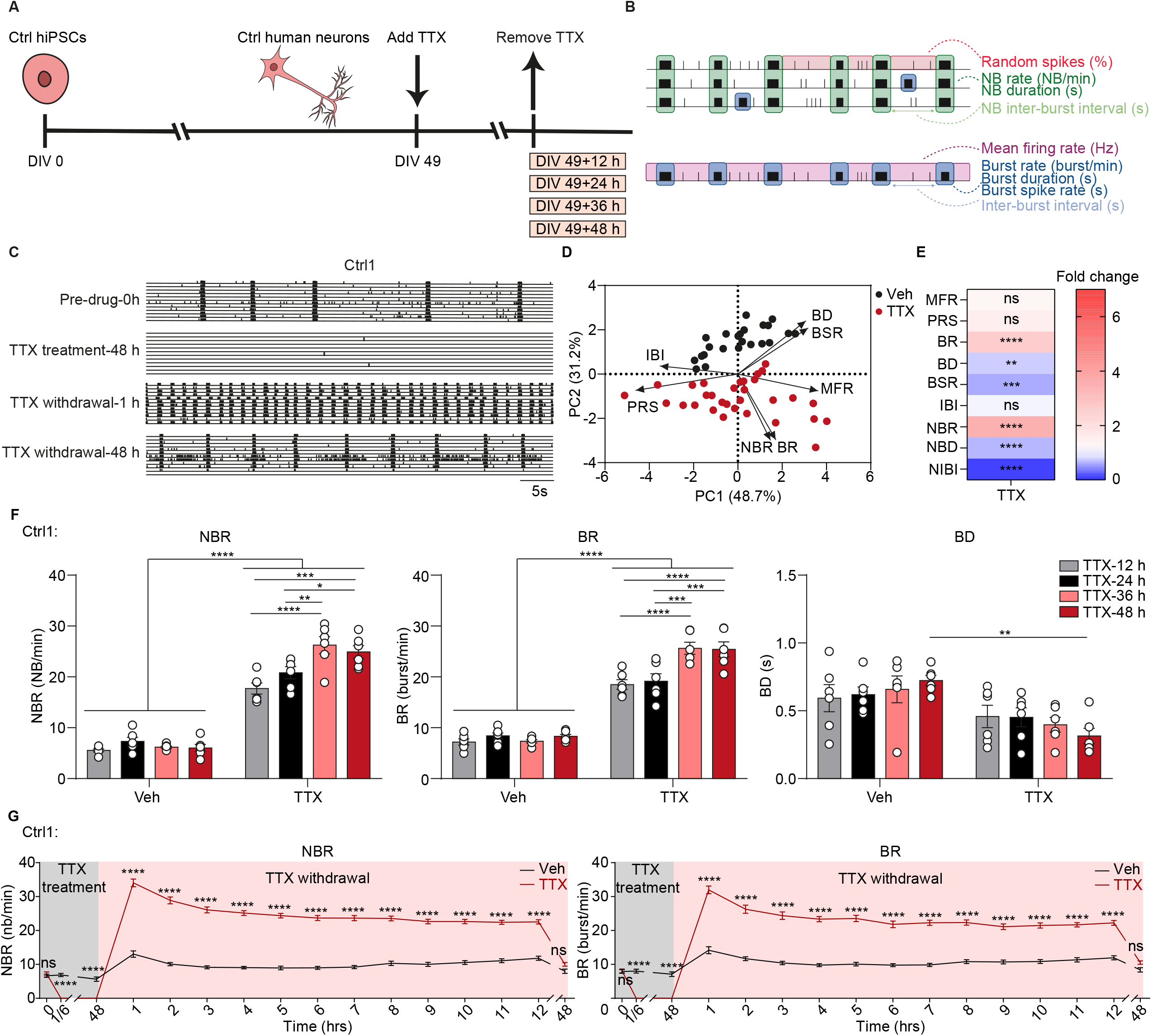
Tetrodotoxin (TTX)-treated neurons show a time-dependent re-arrangement of the networks. (A) Schematic representation of human induced pluripotent stem cell (hiPSC) differentiation and TTX treatment workflow. (B) Schematic overview of extracted parameters from microelectrode arrays (MEAs) recordings. NB = network burst. (C) Representative raster plots showing 1 min of spontaneous activity from hiPSC-derived neuronal networks (Ctrl1) before and after 1 µM TTX treatment, including before addition of TTX (Pre-drug-0 h), 48 h after addition of TTX (TTX treatment-48 h), 1 h after TTX withdrawal (TTX withdrawal-1 h), and 48 h after TTX withdrawal (TTX withdrawal-48 h). (D) Principal component analysis (PCA) plot displaying PC1 and PC2 of all nine analysed MEA parameters for vehicle-treated (Veh) and 48 h TTX-treated (TTX) neurons (Ctrl1 and Ctrl2). All MEA parameters were measured 1 h after TTX withdrawal. MFR = mean firing rate, PRS = percentage of random spike, BR = mean burst rate, BD = mean burst duration, BSR = burst spike rate, IBI = inter-burst interval, NBR = network burst rate, NBD = network burst duration, NIBI = Network burst IBI. n = 23 wells for Veh, n = 31 wells for TTX. (E) Heatmap showing fold changes of all nine analysed MEA parameters after TTX treatment (Ctrl1 and Ctrl2). All MEA parameters were measured at 1 h after TTX withdrawal. n = 23 wells for Veh, n = 31 wells for TTX. (F) Bar graphs showing the effect of 12 h, 24 h, 36 h, and 48 h TTX treatment on the NBR, BR, and BD for Ctrl1 neuronal networks. All MEA parameters were measured 1 h after TTX withdrawal. n = 6 wells for all vehicle-treated (Veh) and TTX-treated (TTX) conditions. (G) Quantification of NBR and BR over time for vehicle-treated (Veh) and 48 h TTX-treated (TTX) neurons (Ctrl1). n = 15 wells for Veh, n = 21 wells for TTX. Data represent means ± SEM. ns: not significant, *P < 0.05, **P < 0.005, ***P < 0.0005, ****P < 0.0001. For panel E, unpaired Student’s T-test was performed between two groups. For panel F and G, two-way ANOVA test followed by a *post-hoc* Bonferroni correction was performed between conditions. All means, SEM and test statistics are listed in Table S3.

TTX treatment for 48 hours (h) on DIV 49 networks completely abolished neuronal activity (Figure 1C). Removing TTX after the 48 h treatment led to a significant increase in network bursting activity, compared with pre-drug and vehicle-treated control conditions (Figure 1C, S1B, and S1E-F). To obtain a full picture of the network changes, we performed a principal component analysis (PCA), including nine independent MEA parameters (Mossink et al., 2021). PCA indicated a clear separation between TTX- and vehicle-treated neuronal networks (Figure 1D). The analysis identified mean network burst rate (NBR), mean burst rate (BR), mean burst duration (BD), and burst spike rate (BSR) as the primary parameters responsible for the observed changes between the two treatment groups (Figure 1D). NBR and BR were increased, while BD, BSR, mean network burst duration (NBD), and network inter-burst interval (NIBI) were decreased following TTX exposure compared with the vehicle-treated condition. In contrast, we found no change in mean firing rate (MFR), percentage of random spike (PRS), and inter-burst interval (IBI) after TTX exposure (Figure 1E). Together, these results suggest that the induction of homeostatic plasticity by prolonged TTX exposure leads to re-arrangement of neuronal networks, without an increase in network activity (as measured by MFR).

Previous studies in rodents have shown that homeostatic plasticity response to TTX at the single-cell level is dependent on the duration of TTX treatment (Benevento et al., 2016; Ibata et al., 2008). We tested if this was also the case at the network level in human neuronal networks by applying TTX for 12, 24, 36, and 48 h. Following the withdrawal of TTX, we observed a significant increase in both NBR and BR already after only 12 h of treatment. These parameters continued to increase with longer treatment durations, peaking at 36 to 48 h of TTX treatment before reaching a plateau. However, BD only decreased after 48 h of TTX exposure, and not with shorter TTX exposures (Figure 1F and S1C). Of interest, when we recorded the network activity on an hourly basis to investigate if the changes in network activity persisted after TTX withdrawal, we found that 48 h TTX-treated neuronal networks gradually returned to pre-drug levels within 48 h (Figure 1G and S1D). This confirms the continuous nature of homeostatic plasticity in human neuronal networks.

### GluA2-lacking AMPA receptors are expressed in an early stage of homeostatic plasticity in hiPSC-derived neurons

Membrane insertion of GluA2-lacking AMPA receptors is required in the early stage of TTX-induced homeostatic plasticity. GluA2-lacking AMPA receptors are subsequently replaced by GluA2-containing AMPA receptors. These mechanisms regulate synaptic strength and are known to underlie synaptic scaling mechanisms (Man, 2011; Soares et al., 2013). To investigate if these mechanisms are also contributing to homeostatic plasticity in this human model system, we applied 1-naphthyl acetyl spermine trihydrochloride (NASPM), a specific antagonist for GluA2-lacking AMPA receptors, to hiPSC-derived neuronal networks that were already exposed to TTX for 12, 24, and 48 h (Figure 2A). We found that NASPM prevented the TTX-induced increase in NBR and BR (Figure 2B-2C) in the neuronal networks treated for 12 h with TTX. Importantly, we corroborated that NASPM had no effect on NBR and BR before TTX application (Figure S2A-2B), confirming that the expression of GluA2-lacking AMPA receptors was exclusively induced by neuronal activity suppression with TTX. After longer TTX exposure, i.e. 24 h and 48 h, NASPM did not affect the increase in NBR and BR (Figure 2D-2G), suggesting that GluA2-lacking AMPA receptors had already been replaced by GluA2-containing AMPA receptors at these time points. Taken together, these results indicate that GluA2-lacking AMPA receptors do contribute to network re-arrangement in hiPSC-derived neuronal networks, where they are incorporated into synapses in an early stage of TTX-induced homeostatic plasticity.

**Figure 2.**
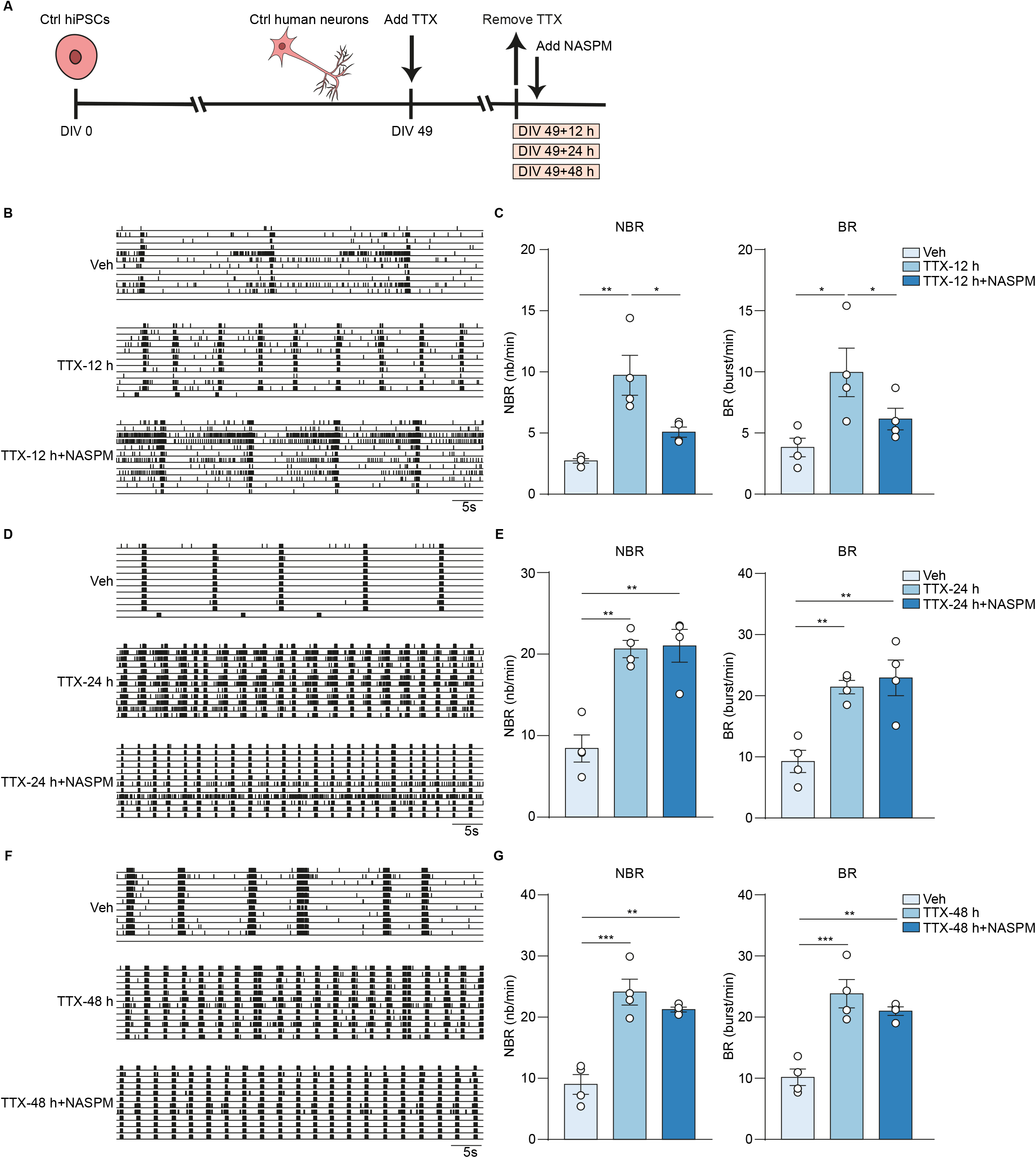
GluA2-Lacking AMPA receptors are expressed in an early stage of homeostatic plasticity. (A) Schematic representation of tetrodotoxin (TTX) treatment and 1-naphthyl acetyl spermine trihydrochloride (NASPM) treatment workflow. (B) Representative raster plots showing 1 min of spontaneous activity from Ctrl2 neuronal networks grown on microelectrode arrays (MEAs). Where indicated, the neurons were vehicle-treated (Veh), or neurons were first treated with 1 µM TTX for 12 h and then TTX was removed (TTX-12 h), or neurons were treated with 10 µM NASPM after TTX was removed (TTX-12 h+NASPM). (C) Bar graphs showing the quantification of mean network burst rate (NBR) and mean burst rate (BR) for (B). n = 4 wells for Veh, n = 4 wells for TTX-12 h, n = 4 wells for TTX-12 h+NASPM. (D) Representative raster plots showing 1 min of spontaneous activity from Ctrl1 neuronal networks grown on MEAs. Where indicated, the neurons were vehicle-treated (Veh), or neurons were first treated with 1 µM TTX for 24 h and then TTX was removed (TTX-24 h), or neurons were treated with 10 µM NASPM after TTX was removed (TTX-24 h+NASPM). (E) Bar graphs showing the quantification of NBR and BR for (D). n = 4 wells for Veh, n = 4 wells for TTX-24 h, n = 4 wells for TTX-24 h+NASPM. (F) Representative raster plots showing 1 min of spontaneous activity from Ctrl1 neuronal networks grown on MEAs. Where indicated, the neurons were vehicle-treated (Veh), or neurons were first treated with 1 µM TTX for 48 h and then TTX was removed (TTX-48 h), or neurons were treated with 10 µM NASPM after TTX was removed (TTX-48 h+NASPM). (G) Bar graphs showing the quantification of NBR and BR for (F). n = 4 wells for Veh, n = 4 wells for TTX-48 h, n = 4 wells for TTX-48 h+NASPM. Data represent means ± SEM. *P < 0.05, **P < 0.005, ***P < 0.0005, one-way ANOVA test followed by a *post-hoc* Bonferroni correction was performed between conditions. All means, SEM and test statistics are listed in Table S3.

### Homeostatic plasticity involves increased miniature excitatory postsynaptic current amplitude and elongation of axon initial segments in hiPSC-derived neurons

We next investigated pre-synaptic and post-synaptic contributions to synaptic scaling following TTX treatment at the single-cell level. With whole-cell electrophysiological recordings of hiPSC-derived neurons treated with vehicle or TTX for 48 h, we found that TTX treatment induced an increase in miniature excitatory postsynaptic current (mEPSC) amplitude without affecting mEPSC frequency (Figure 3A-3C and S3A-3C). The changes in mEPSC amplitude were scalable as they were observed across different mEPSC amplitudes (Figure 3D and S3D), which is a major property of synaptic scaling (Moulin et al., 2020; Turrigiano et al., 1998). This functional readout suggests that neuronal activity suppression results in increased post-synaptic AMPA receptor expression. To corroborate these results, we quantified the expression of surface GluA2-containing AMPA receptors at the post-synaptic membrane after TTX treatment. Analysis of GluA2 surface expression indeed revealed an increase in the number of surface GluA2 puncta after 48 h TTX treatment (Figure 3E-3F), which is in line with our functional data at single-cell and neuronal network levels.

**Figure 3.**
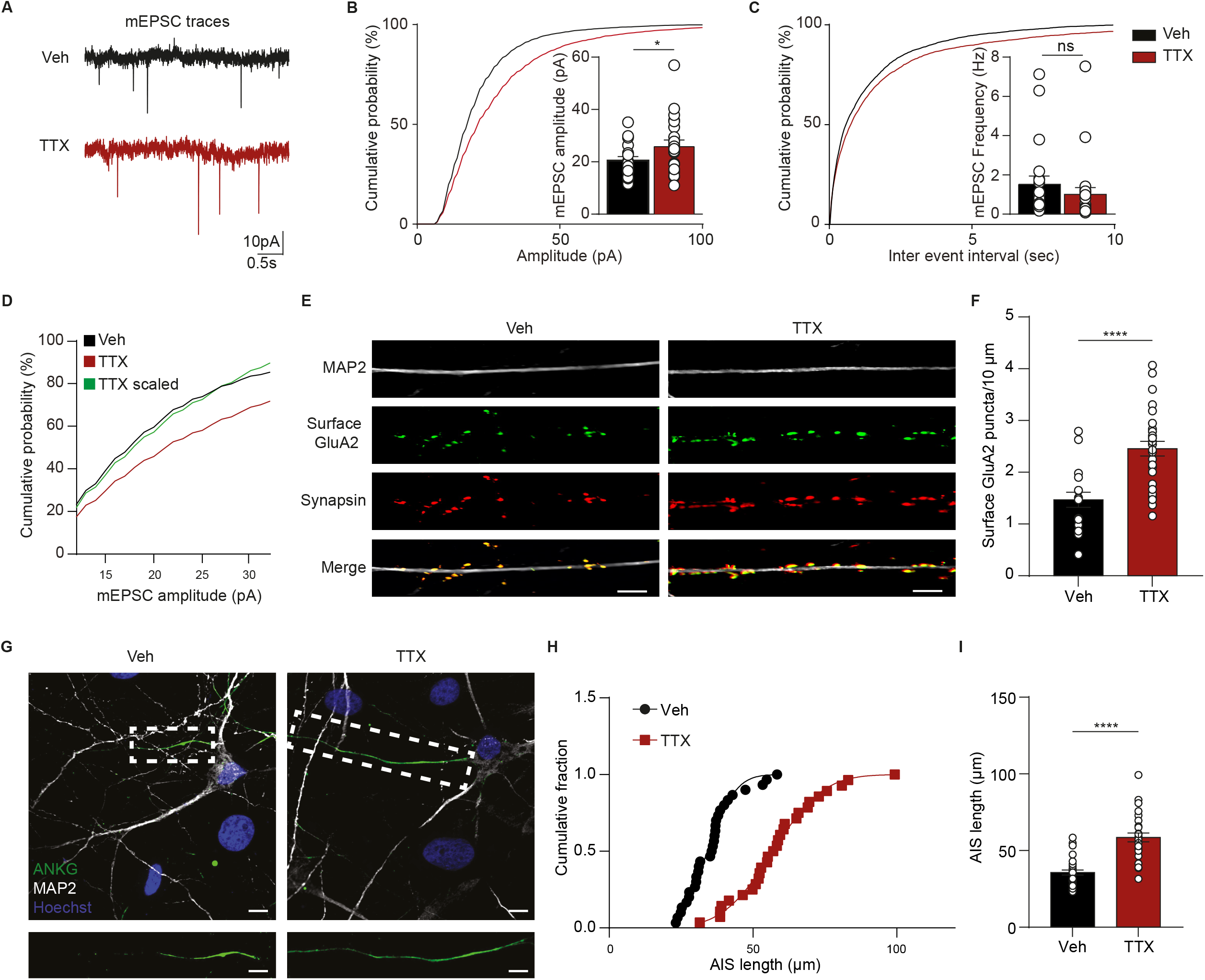
Increased miniature excitatory postsynaptic current (mEPSC) amplitudes and elongation of axon initial segment (AIS) following TTX treatment. (A-D) Representative mEPSC traces of vehicle-treated (Veh) and 48 h TTX-treated (TTX) neurons (Ctrl1; A). Quantification of the amplitude (B) and frequency of mEPSCs (C) in Veh- and TTX-treated neurons. n = 21 recordings for Veh, n = 23 recordings for TTX (in total 3 batches for each condition). Rescaled cumulative mEPSCs amplitude (Scaling = TTX values * 1.21; D). (E) Representative images of vehicle-treated (Veh) and 48 h TTX-treated (TTX) neurons (Ctrl1) stained for MAP2 (grey), surface GluA2 (green), and synapsin 1/2 (red) (scale bar 5 µm). (F) Quantification of GluA2 puncta (number per 10 µm). n = 18 cells for Veh, n = 28 cells for TTX. (G) Representative images of vehicle-treated (Veh) and 48 h TTX-treated (TTX) neurons (Ctrl1) stained for MAP2 (grey), ANKG (green), and Hoechst (blue) at DIV 49 (scale bar 5 µm). (H-I) Plots showing cumulative fraction and quantification of the length axon initial segment (AIS; µm). n = 14 cells for Veh, n = 14 cells for TTX. Data represent means ± SEM. ns: not significant, *P < 0.05, ****P < 0.0001, unpaired Student’s T-test was performed between two groups. All means, SEM and test statistics are listed in Table S3.

In addition to synaptic scaling, neurons also respond to altered activity by modifying their intrinsic excitability. Changes in neuronal intrinsic excitability can occur through a variety of mechanisms, including changes in structural characteristics of the axon initial segment (AIS). For instance, the elongation of the AIS has been shown to increase neuronal excitability and to facilitate action potential (AP) generation (Evans et al., 2015; Li et al., 2020). TTX treatment (48 h) significantly increased AIS length compared to vehicle-treated neurons (Figure 3G-3I). This confirms that, beside changes in functional synaptic properties, alterations in the structural characteristics of axons also occur following neuronal activity suppression with TTX and may contribute to the increase in network excitability.

### Transcriptional changes in neurons following TTX treatment are associated with homeostatic plasticity

The timeframe of homeostatic plasticity-related processes suggests that changes in the transcription program of neurons could mediate such processes (Ibata et al., 2008; Schaukowitch et al., 2017). To study the potential changes in gene expression activity induced by neuronal activity suppression, in both neurons and astrocytes, we performed RNA-sequencing (RNA-seq) of vehicle- and 48 h TTX-treated DIV 49 neuron-astrocyte co-cultures. PCA analysis segregated TTX- and vehicle-treated neurons and astrocytes, confirming TTX treatment-related changes in the transcriptional activity of the neuronal networks (Figure 4A and 5A).

**Figure 4.**
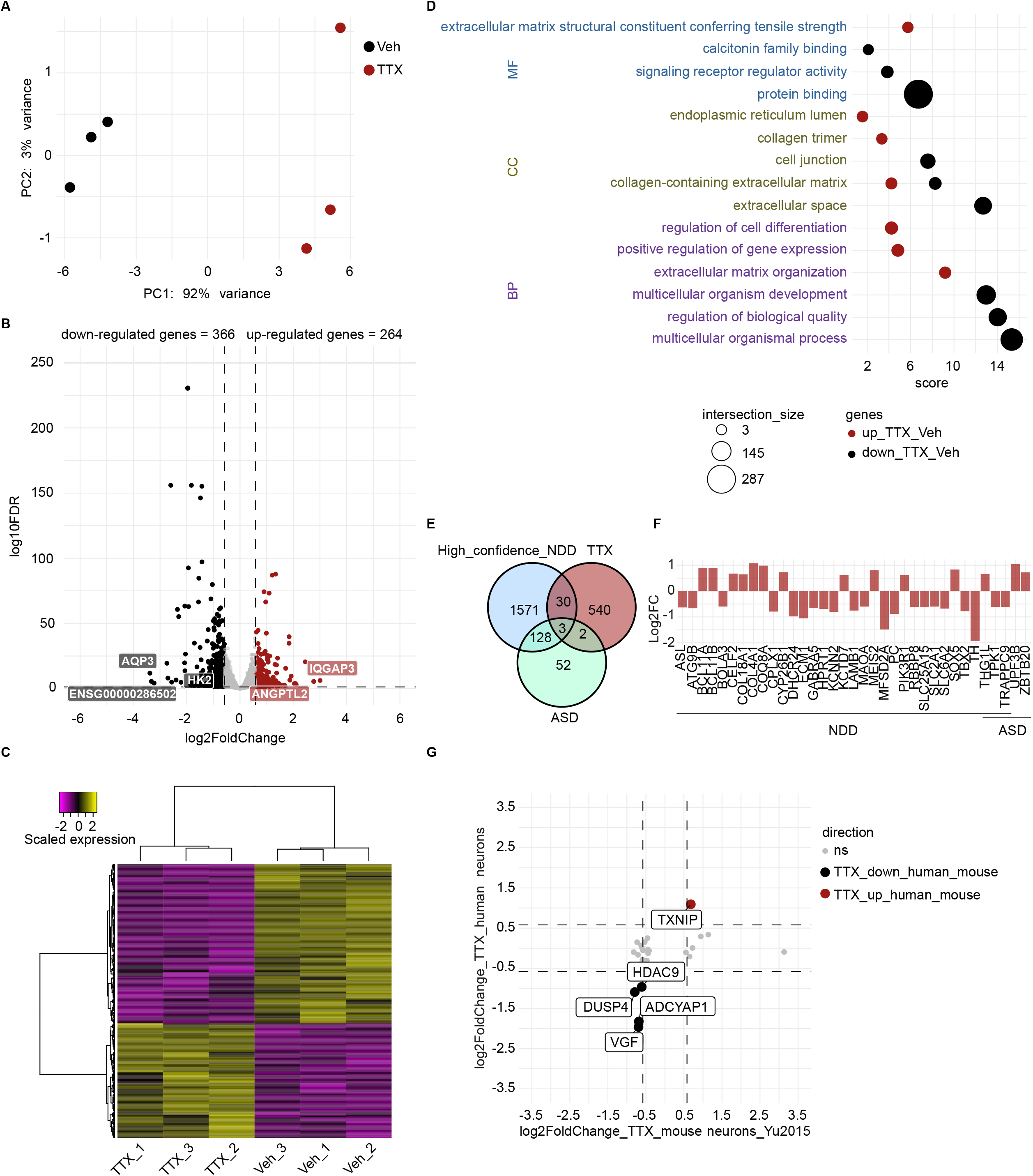
Transcriptional changes in human neurons associated with homeostatic plasticity. (A) Principal component analysis (PCA) of RNA-sequencing data of six samples, three TTX- and three vehicle-treated hiPSC-derived neurons samples (Ctrl1). Colours indicate the received treatment (red=TTX, and black=vehicle). (B) Volcano plot depicting differential gene expression between TTX- and vehicle-treated neurons. Coloured dots indicate differentially expressed genes (DEGs; absolute log_2_ fold change (Log_2_FC) > 0.58 and adjusted P value < 0.05). Up-regulated genes in TTX-treated neurons are shown in red, down-regulated genes are depicted in black. The five genes with the greatest change in expression activity between the two conditions are indicated. (C) Heatmap depicting scaled expression of DEGs in the six samples. (D) Scatter plot of the enrichment analysis results depicting the top 3 significant gene ontology (GO) terms associated with up-regulated (red dots) and down-regulated genes (black dots) in neurons exposed to TTX treatment as compared to vehicle-treated neurons. Terms are ordered per ontological category (MF: molecular function; CC: cellular component; BP: biological process). The size of the dots indicates the number of genes observed in the list of DEGs and the list of genes associated with the particular term. Score: negative log_10_ of the adjusted P value resulting from the enrichment analysis. (E) Venn diagram depicting the number of shared genes between the list of DEGs in TTX-treated neurons (red circle, TTX) and genes previously related to neurodevelopmental disorders (NDDs; light-blue circle, High_confidence_NDD) (Leblond et al., 2021) and highly confident autism-related genes (light-green circle, ASD) (Fu et al., 2022). (F) Bar plot depicting the Log_2_FC value of the 35 shared genes observed in (E). Fold change was calculated by normalizing gene expression level of TTX-treated condition to vehicle-treated condition. (G) Four-way volcano plot depicting expression changes in hiPSC-derived neurons and mouse neurons treated with TTX (absolute Log_2_FC > 0.58 and adjusted P value < 0.05) (Yu et al., 2015). Each dot represents a gene. Red dots represent genes up-regulated in both hiPSC-derived neurons and mouse neurons in response to TTX, black dots represent down-regulated genes in both hiPSC-derived neurons and mouse neurons exposed to TTX.

**Figure 5.**
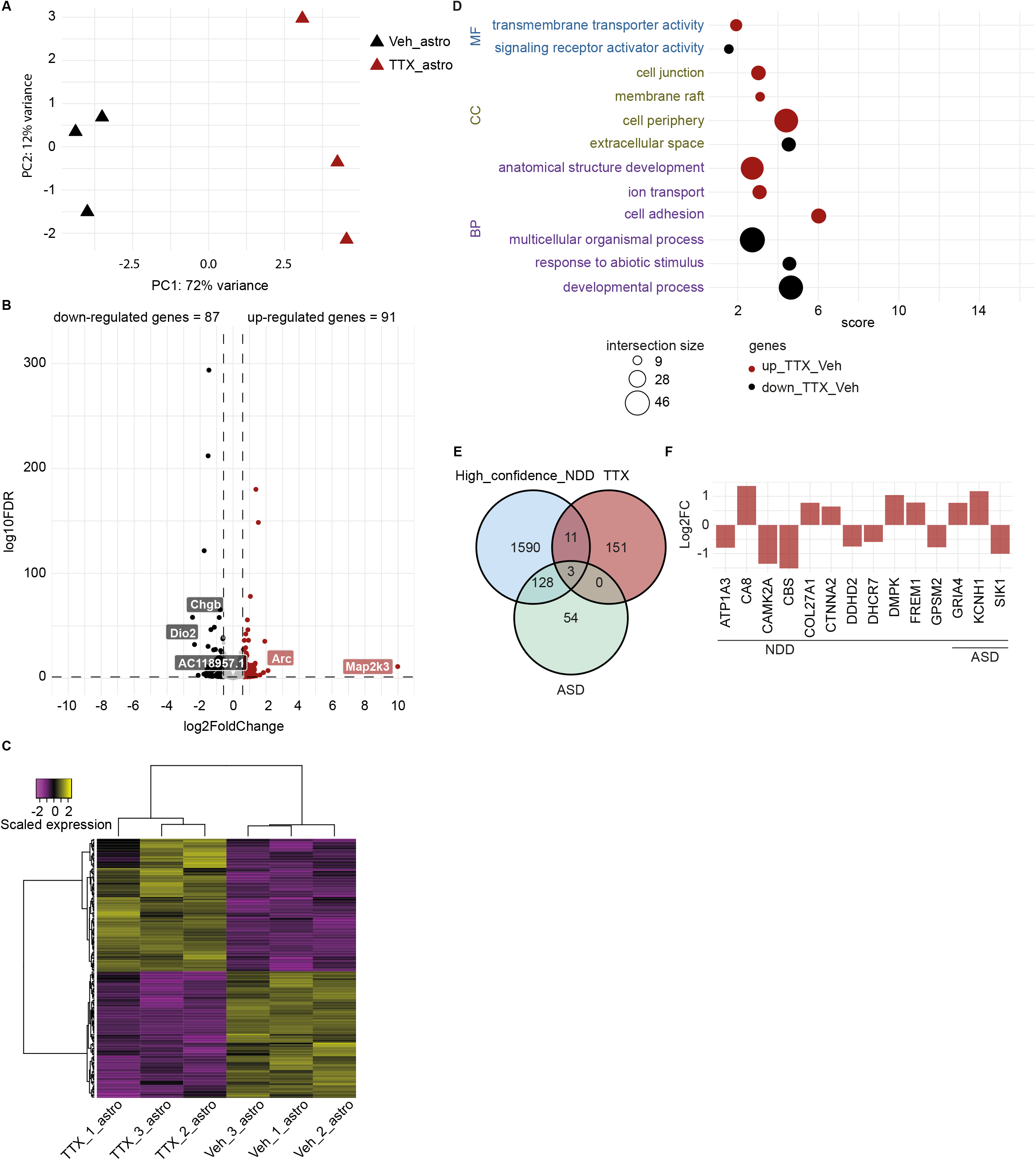
Transcriptional changes are induced by suppression of neuronal activity in rat astrocytes. (A) PCA plot of RNA-sequencing data of six samples, three TTX- and three vehicle-exposed rat astrocytes derived from co-cultures with hiPSC-derived neurons (Ctrl1). Colours indicate the received treatment (red=TTX, and black=vehicle). (B) Volcano plot depicting differential gene expression between TTX- and vehicle-exposed astrocytes. Coloured dots indicate DEGs (Absolute Log_2_FC > 0.58 and adjusted P value < 0.05). Up-regulated genes in TTX-exposed astrocytes are shown in red, down-regulated genes are depicted in black. The five genes with the greatest change in expression activity between the two conditions are indicated. (C) Heatmap depicting scaled expression of DEGs in the six astrocyte samples. (D) Scatter plot depicting the top 3 significant GO terms, per ontological category (molecular function: MF, cellular component: CC and biological process: BP), associated with up-regulated (red dots) and down-regulated genes (black dots) in astrocytes exposed to TTX treatment, as compared to vehicle-exposed astrocytes. The size of the dots indicates the number of intersected genes between the list of DEGs and the genes associated with the particular term. Score: negative logarithm10 of the adjusted P value resulting from the enrichment analysis. (E) Venn diagram depicting the number of shared genes between the list of DEGs in TTX-exposed astrocytes (red circle, TTX) and genes previously related to NDDs (light-blue circle, High_confidence_NDD) (Leblond et al., 2021) and highly confident autism-related genes (light-green circle, ASD) (Fu et al., 2022). (F) Bar plot depicting the Log_2_FC value of the 14 shared genes observed in (E). Fold change was calculated by normalizing gene expression level of TTX-treated condition to vehicle-treated condition.

With an absolute log_2_ fold change (FC) > 0.58 and multiple testing-adjusted P value < 0.05, we identified 366 down-regulated and 264 up-regulated genes in TTX-treated neuronal networks, compared to vehicle-treated ones (Figure 4B; Table S1). These transcriptional changes were consistent across samples within each condition (Figure 4C). The largest changes in gene expression were observed for *IQGAP3*, *ANGPTL2*, *AQP3*, *ENSG00000286502*, and *HK2* (Figure 4B; Table S1).

Among genes up-regulated by TTX treatment, gene ontology (GO) analysis showed “extracellular matrix organization” to be the most strongly enriched term, including members of the Disintegrin and Metalloproteinase with Thrombospondin motifs (ADAMTS) family, such as *ADAMTS17*, *ADAMTS8*, and *ADAMTS18* (Figure 4D and S4A; Table S1). Additionally, the term “positive regulation of gene expression” was also enriched, including several transcription factors, such as *ATF3* and *HOXA5* (Figure S4A; Table S1).

Genes down-regulated by TTX-treatment, including *ADAMTSL4*, *ADAMTS9, NPTX1, BDNF, VGF, NR4A1,* and *TNC* (Figure S4A; Table S1), were generally related to cell communication; enriched terms included “response to stimulus”, “signalling”, “transport”, “vesicle”, “extracellular space”, and “cell junction” (Figure 4D; Table S1). Neuronal pentraxin-1 (NPTX1) and brain-derived neurotrophic factor (BDNF) have been previously implicated in homeostatic plasticity (Schaukowitch et al., 2017; Fernandes and Carvalho, 2016), and we corroborated the reduction of *BDNF* and *NPTX1* expression in the TTX-treated networks with quantitative polymerase chain reaction (Figure S4B). By assessing the function of TTX-induced differentially expressed genes (DEGs) in the synaptic compartment through SynGO ontologies and annotations (Koopmans et al., 2019), we observed that DEGs were mostly associated with synaptic organization, synaptic signaling, presynaptic and postsynaptic processes (Figure S4C and S4D; Table S1).

Considering that alterations in homeostatic plasticity during development may contribute to the pathophysiology of different neurodevelopmental disorders (NDDs; Antoine et al., 2019; Bulow et al., 2019; Lee et al., 2018; Benevento et al., 2016; Zeng et al., 2007; Dani et al., 2005), we tested whether the TTX-induced transcriptional changes were associated with genes previously related to NDDs (Fu et al., 2022; Leblond et al., 2021). While we did not identify a significant association between the list of TTX-induced DEGs and lists of genes previously related to NDDs (Table S1), we still observed some overlap (5.6%; 35 of 630; Figure 4E and 4F), including *MAOA, TH,* and *TRAPPC9* (Figure 4F; Table S1). This result suggests that altered expression of these genes could affect homeostatic plasticity in some NDDs.

Finally, to identify potential conserved molecular mechanisms involved in homeostatic plasticity between human and mouse neurons, we compared our list of TTX-induced DEGs with previously identified TTX-induced DEGs in mouse primary hippocampal neurons (Yu et al., 2015). The overlap with previously identified up-regulated (0.4%; 1 of 264) and down-regulated DEGs was however low (1%; 4 of 366; Figure 4G; Table S1). Overlapping genes were *TXNIP, HDAC9*, *DUSP4*, *ADCYAP1*, and *VGF* (Figure 4G; Table S1). This may suggest that changes in gene expression activity contributing to homeostatic plasticity differ between mice and humans, emphasizing the importance of using human models to investigate homeostatic plasticity in order to gain a more accurate understanding of how it works in humans specifically.

Overall, these results identify multiple genes that likely contribute to homeostatic plasticity. Suppression of neuronal activity may induce reduction in the expression of genes involved in cell communication, while changes in the extracellular matrix might contribute to increased neuronal synchronized activity observed in homeostatic plasticity.

### Suppression of neuronal activity induces transcriptional changes in astrocytes

Astrocytes are sensitive to changes in neuronal activity and are known regulators of homeostatic plasticity (Lines et al., 2020; Perez-Catalan et al., 2021). We identified 91 up-regulated and 87 down-regulated astrocytic genes in response to neuronal activity suppression with TTX (Figure 5B; Table S2). Transcriptional changes in astrocytes were consistently seen in same conditions (Figure 5C; Table S2). *Arc*, *Map2k3*, *Dio2*, *Chgb*, and *AC118957.1* were identified as the genes with the strongest expression changes between vehicle- and TTX-exposed astrocytes (Figure 5B; Table S2).

GO enrichment analysis indicated significant association between the up-regulated astrocytic genes and ontological terms likely related to neuronal function (Figure 5D; Table S2), such as “cell junction”, “ion transport”, “neuron projection”, and “transmembrane transporter activity”. Particularly, the expression of *Arc*, *Egr3, Avpr1a*, and *Astn2* was increased in TTX-exposed astrocytes (Figure S5A; Table S2). Together, these observations suggest that upon suppression of neuronal activity, astrocytes increase the expression of genes that could modulate synaptic activity. In contrast, down-regulated genes in astrocytes exposed to TTX showed significant association with terms related to “response to abiotic stimulus”, “response to external stimulus”, and “signalling receptor activity”, suggesting that a reduced expression of genes related to response to stimuli in astrocytes may be a consequence of neuronal activity suppression, which could decrease the stimulation of the astrocytes in culture. We further assessed the role of astrocytic DEGs induced by neuronal activity suppression in the synaptic compartment using SynGO ontologies and annotations (Koopmans et al., 2019); we observed that astrocytic DEGs were related to both presynaptic and postsynaptic processes (Figure S5C and S5D; Table S2).

Similar to our analysis for the hiPSC-derived neurons, we compared the list of DEGs in astrocytes exposed to neuronal activity suppression with lists of genes previously related to NDDs (Fu et al., 2022; Leblond et al., 2021). We found that 7.9% (14 of 178) of DEGs were known NDD genes (Figure 5E; Table S2), such as *Camk2a* (Figure 5F; Table S2). Their observed roles in NDDs might thus be related to an impaired astrocytic contribution to homeostatic plasticity.

Finally, to identify potentially shared mechanisms between neurons and astrocytes that could contribute to homeostatic plasticity in our co-cultures, we compared the transcriptional changes induced by TTX treatment in the two cell types. To this end, we converted rat gene symbols to human gene symbols and combined the transcriptional profiles of all sequenced samples, including astrocytes and neurons. PCA showed cell type to be the major source of variation in gene expression. Nonetheless, samples were also segregated by treatment, suggesting some shared transcriptional changes in neurons and astrocytes after neuronal activity suppression (Figure S5B). When intersecting the list of DEGs by TTX treatment in neurons and astrocytes, we identified 11 commonly down-regulated genes, including *VEGFA*, *SCG2*, and *CCK*, while the expression of *ATF3* and *COL13A1* was increased in both cell types. We also identified 9 genes with opposite changes in transcriptional activity between neurons and astrocytes, including *SPON1*, *IGFBP3*, and *DISP3* (Figure S5E). These oppositely regulated genes, of which several are involved in lipid and energy metabolism, may be involved in crosstalk mechanisms between neurons and astrocytes that contribute to homeostatic plasticity.

Concluding, our transcriptional analysis of astrocytes in the neuronal networks provides evidence corroborating their role in the regulation of homeostatic plasticity and identifies candidates for underlying genes and biological pathways.

## Discussion

In this study, we describe a human *in vitro* neuronal model for studying homeostatic plasticity at the network level. Using this model, we provide insight into how synaptic strength and intrinsic properties of neurons cooperate to stabilize neuronal network activity in response to activity suppression. We demonstrated that chronic deprivation of neuronal activity through the inhibition of voltage-gated sodium channels with TTX elicited a time-dependent re-arrangement of neuronal networks, reflected in increased synchronized network activity. TTX-induced modifications in neuronal network properties were accompanied by increased surface expression of post-synaptic AMPA receptors, as well as by increased neuronal excitability. Additionally, we identified transcriptional changes induced by suppression of neuronal activity in neurons and astrocytes, which may underlie the network re-arrangement. At the network level, increased synchronized activity following neuronal activity suppression has been described in a rodent model of homeostatic plasticity, where short-term treatment with TTX induced synchronized burst oscillations (Zhou et al., 2009). Our study in hiPSC-derived neurons verified earlier reported presence of homeostatic plasticity mechanisms regulating neuronal excitability at the network and single-cell level (Cordella et al., 2022; Zhang et al., 2018). We found that in the hiPSC-derived neurons, in an early stage, the increase in synchronized network activity observed after suppression of neuronal activity was dependent on the expression of GluA2-lacking AMPA receptors and, in a later stage, on the expression of GluA2-containing AMPA receptors. This is consistent with a previous study in rodents showing a role for GluA2-lacking AMPA receptors in the induction of homeostatic plasticity (Man, 2011). Moreover, we confirmed that the network re-arrangement following TTX treatment was associated with increased mEPSC amplitudes and elongation of the AIS, matching expected changes in structural properties of AIS and postsynaptic AMPA receptors after suppression of neuronal activity (Chater and Goda, 2014; Grubb and Burrone, 2010). Taken together, these results indicate that we provide a valid model of homeostatic plasticity. Using our model system, we observed that increased synchronized activity gradually returned to pre-drug level within 48 h after TTX removal, consistent with the notion that homeostatic plasticity, as induced by TTX, will restore network activity from a hyper-active level to a set-point level. In mammalian synapses, pharmacological block of postsynaptic AMPA receptors or genetic knock-out of GluA4 AMPA receptors is known to trigger a presynaptic form of homeostatic plasticity (Delvendahl et al., 2019; Jakawich et al., 2010; Lindskog et al., 2010). Thus in our model, we may hypothesize that the accumulation of post-synaptic AMPA receptors, as induced by the prolonged TTX exposure, might cause an increase in miniature excitatory postsynaptic potential amplitude, which then induces a homeostatic decrease in presynaptic vesicle release. Future studies are needed to test this hypothesis.

Previous studies in rodents suggested that transcriptional changes contribute to homeostatic plasticity (Schaukowitch et al., 2017; Benevento et al., 2016; Yu et al., 2015). In our study, the transcriptional program induced by neuronal activity suppression implicated increased expression of neuronal genes related to “extracellular matrix organization”. Extracellular matrix (ECM) molecules are synthesized by neurons as well as by non-neural cells, and are secreted into the extracellular space (Dityatev and Schachner, 2003). Re-organization of the extracellular matrix can modulate neuronal connectivity and plays a role in synaptic plasticity and homeostasis (Bikbaev et al., 2015; Frischknecht and Gundelfinger, 2012; Dityatev et al., 2010). Members of the ADAMTS family, such as *ADAMTS17*, *ADAMTS8*, and *ADAMTS18*, exhibited increased expression in response to TTX (Figure S4A). These proteins are involved in the degradation of the ECM during development, and play an essential role in neuroplasticity (Ferrer-Ferrer and Dityatev, 2018; Gottschall and Howell, 2015). Based on our results, the reorganization of the ECM also appears important in homeostatic plasticity. Other ECM molecules also were altered in their expression by suppression of neuronal activity, such as tenascin C (TNC), which exhibited reduced neuronal expression in response to TTX (Figure S4A). Reduced TNC might cause the increase of mEPSC amplitude we observed by reducing L-type voltage-gated calcium channel-mediated signaling (Evers et al., 2002; Wang et al., 2011). Genes involved in “positive regulation of gene expression”, were up-regulated in the neurons after TTX treatment, including transcription factors, such as *ATF3* and *HOXA5* (Figure S4A). These transcription factors have previously been shown to regulate synaptic transmission and synaptic plasticity (Ahlgren et al., 2014; Lizen et al., 2017), and they may be key modulators of the homeostatic responses. Conversely, down-regulated neuronal genes were related to cell communication in response to TTX-treatment. This could be a mere consequence of suppressing neuronal activity and may not be necessarily linked directly to molecular mechanisms underlying homeostatic plasticity. For instance, the expression of *NR4A1* is activity-dependent (Jeanneteau et al., 2018; Hawk and Abel, 2011), and we observed reduced *NR4A1* in TTX-treated neurons. Feedback loops may be hypothesized, by which the detection of a reduced expression of genes related to cell communication could contribute to activating homeostatic plasticity mechanisms.

Our *in vitro* model of network plasticity allows us to explore not only neuronal mechanisms of homeostatic plasticity but also the potential role of astrocytes in this process. In our study, we observed significant transcriptional changes in astrocytes exposed to neuronal activity suppression. For example, *Arc* (Figure S5A), an immediate-early gene (IEG), was upregulated in its expression. In neurons, Arc expression is dynamically regulated by neuronal activity, and it modulates synaptic strength by controlling endocytosis of AMPA receptors at the synapse (Shepherd and Bear, 2011; Shepherd et al., 2006). It has previously been reported that *Arc* and other IEGs are also expressed in astrocytes (Rodriguez et al., 2008; Rubio, 1997; Kato et al., 1995). We speculate that *Arc* was transcribed and may be released from the astrocytes into the synapse, where it may modulate synaptic plasticity, as described for other astrocyte-secreted factors (Perez-Catalan et al., 2021; Wang et al., 2021). We also observed increased expression of *Egr3, Avpr1a*, and *Astn2* in TTX-exposed astrocytes (Figure S5A). Avpr1a belongs to the subfamily of G-protein-coupled receptors that bind arginine vasopressin (AVP), and it promotes long-term potentiation (LTP) induction (Namba et al., 2016); Egr3 is important for normal hippocampal long-term depression (LTD) (Gallitano-Mendel et al., 2007); Astn2 is a transmembrane protein that also modulates synaptic function by trafficking of neuroligins and synaptic proteins in Purkinje cells (Behesti et al., 2018). Together, these observations suggest that astrocytes increase the expression of genes that could modulate synaptic activity upon suppression of activity of (neighboring) neurons. Such findings indicate that the astrocytic contribution to homeostatic plasticity might be more pronounced than previously considered and should be studied more intensively.

Although not statistically over-represented, several genes differentially expressed by TTX-treatment have previously been associated with increased risk of neurodevelopmental disorders (NDDs) through genetic mutation, including *KCNN2* and *ZBTB20*. *KCNN2* encodes a voltage-independent potassium channel activated by intracellular calcium, and is thought to regulate neuronal excitability and synaptic transmission (Willis et al., 2017; Adelman et al., 2012; Ballesteros-Merino et al., 2012; Lin et al., 2008). The role of KCNN2 in homeostatic plasticity, through dysregulation of neuronal excitability (Raghuram et al., 2017), may explain its link to NDDs. Similarly, *ZBTB20*, which encodes a zinc finger protein with an essential role in LTP and memory formation through modulation of NMDA receptor activity and activation of ERK and CREB (Ren et al., 2012), may be linked to NDDs also through its role in homeostatic plasticity. In astrocytes, the NDD-linked gene *Camk2a* (Kury et al., 2017) was differentially expressed after neuronal activity suppression with TTX. *Camk2a* encodes a calcium calmodulin-dependent protein kinase II Alpha (CaMKII; Silva et al., 1992). Inhibition of CaMKII signaling in cortical astrocytes reduces glutamate uptake and induces neurotoxic release of ATP (Ashpole et al., 2013). NDD-linked genetic mutations in *Camk2a* might act through an altered astrocytic contribution to homeostatic plasticity, as glutamate uptake plays a critical role in regulating the strength and the extent of receptor activation at excitatory synapses (Valtcheva and Venance, 2019). Taken together, our data highlights that mechanisms of neurodevelopmental disorders may involve homeostatic plasticity-related gene functions.

Some limitations should be taken into account in the interpretation of our results. Considering that some sub-types of astrocytes exhibit sensitivity to TTX (McNeill et al., 2021; Sontheimer and Waxman, 1992), we cannot discriminate transcriptional changes directly related to suppression of neuronal activity and homeostatic plasticity mechanisms, from changes induced by TTX effects on astrocytes (that may or may not influence homeostatic plasticity). Comparing the gene expression profile of our TTX-treated neuron-astrocyte co-cultures with pure astrocyte cultures treated with TTX may help identify genes and processes induced by TTX treatment in astrocytes and provide a basis for testing their role in homeostatic plasticity. Also, the application of other methods to induce homeostatic plasticity, more complex co-cultures, and setups using human astrocytes will improve the predictive value of our model.

To conclude, we provide evidence that hiPSC-derived neuronal networks display homeostatic plasticity at the network and single-cell level. Our *in vitro* model of network plasticity is versatile and allows investigation of human homeostatic plasticity mechanisms. It may also provide a platform for (high-throughput) drug screening, to explore whether specific compounds can be used to rescue altered or insufficient homoeostatic plasticity in the context of brain disorders, such as NDDs.

## Experimental procedures

### Human iPSCs cell culture

In this study, we used three independent control human induced pluripotent stem cell (hiPSC) lines, including Ctrl1, Ctrl2, and Ctrl3. All of them were obtained by reprogramming skin fibroblasts and have been described in detail previously (Frega et al., 2019; Mossink et al., 2021). More details also can be found in our supplemental experimental procedures. HiPSCs were cultured in Essential 8 Flex medium (Gibco, A2858501) supplemented with 0.1 mg/mL Primocin (Invivogen, ant-pm-2) on 6-well plates pre-coated with Matrigel (1:15 diluted in DMEM/F12 medium; Matrigel: Corning, #356231; DMEM/F12 medium: Gibco, 11320074) at 37°C, 5% CO_2_. HiPSCs were infected with Neurogenin-2 (Ngn2) and rtTA lentivirus. The vector utilized for generation of the rtTA lentivirus was pLVX-EF1α-(Tet-On-Advanced)-IRES-G418(R), which encodes a Tet-On advanced trans-activator under control of a constitutive EEF1A/EF1α promoter and has resistance to the antibiotic G418. The lentiviral vector for Ngn2 was pLVX- (TRE-tight)-(MOUSE)Ngn2-PGK-Puromycin(R), encoding Nng2 under control of a Tet-controlled promoter and the puromycin resistance gene under control of a constitutive PGK promoter. Both vectors were being transfected and packaged into lentiviral particles through using the packaging vectors psPAX2 lentiviral packaging vector (Addgene, 12260) and pMD2.G lentiviral packaging vector (Addgene, 12259). Medium was supplemented with puromycin (0.5 g/mL) and G418 (50 g/mL). Medium was refreshed every other day, and cells were passaged 1-2 times per week using an enzyme-free reagent (ReLeSR; Stem Cell Technologies, 05872). Cells were checked for mycoplasma contamination every two weeks. Droplet digital PCR (ddPCR) was performed to test genome integrity.

### Neuronal differentiation

At days *in vitro* (DIV) 0, hiPSCs were dissociated with Accutase (Sigma-Aldrich, A6964) to generate single cells. HiPSCs were then plated on microelectrode array (MEA) plates (20,000 cells/well) or glass, nitric-acid treated coverslips (20,000 cells/well) in Essential 8 medium (Gibco, A1517001) supplemented with Primocin (0.1 mg/mL, Invivogen, ant-pm-2), RevitaCell (1:100, Gibco, A2644501), and doxycycline (4 µg/mL). MEA plates and coverslips were pre-coated with 50 µg/mL poly-L-ornithine (Sigma-Aldrich, P4957) for 3 h at 37°C, 5% CO_2_, and 20 µg/mL human recombinant laminin (BioLamina, LN521) overnight at 4°C. At DIV 1, the medium was changed to DMEM/F12 (Gibco, 11320074), which was supplemented with MEM non-essential amino acid solution NEAA (1:100, Sigma-Aldrich, M7145), N-2 Supplement (1:100, Gibco, 17502048), BDNF (10 ng/mL, PeproTech, 450-02), NT-3 (10 ng/mL, PeproTech, 450-03), doxycycline (4 µg/mL, Sigma, D9891), Primocin (0.1 mg/mL, Invivogen, ant-pm-2), and mouse laminin (0.2 mg/mL, Sigma-Aldrich, L2020). At DIV 2, freshly prepared rat astrocytes were added, in a 1:1 ratio, in order to support neuronal maturation. At DIV 3, the medium was fully changed to Neurobasal medium (Gibco, 21103049), which was supplemented with B-27 Supplement (1:50, Gibco, 17504044), GlutaMAX (1:100, Gibco, 35050038), Primocin (0.1 mg/mL, Invivogen, ant-pm-2), NT-3 (10 ng/mL, PeproTech, 450-03), BDNF (10 ng/mL, PeproTech, 450-02), and doxycycline (4 µg/mL). Cytosine β-D-arabinofuranoside (Ara-C, 2 µM, Sigma-Aldrich, C1768) was added once to remove proliferating cells from the cultures. From DIV 6 onwards, medium was 50% refreshed every other day. From DIV 10 onwards, the medium was additionally supplemented with 2.5% fetal bovine serum (FBS, Sigma, F2442) to support astrocyte viability. Neurons were differentiated to glutamatergic neurons by overexpression of Ngn2 until DIV 30 or DIV 49.

### Procedure for TTX treatment and TTX withdrawal on MEA plates

All experiments were performed at DIV 49, as younger neurons were very sensitive to the washing out process, reflected in overshoot neuronal network activity at DIV 30, but not at DIV 49 (Figure S1A). At DIV 48, 50% pre-conditioned untreated culture medium was collected, and medium was then 50% refreshed. Specifically, 250 µL of the spent medium was collected in a 15 mL tube and stored at 4°C. Of the freshly prepared medium, 250 µL was added to cells. At DIV 49, a first MEA recording was performed by using the 24-well MEA system (Multichannel Systems, MCS GmbH, Reutlingen, Germany). After the MEA recording, 1 μM tetrodotoxin (TTX, Tocris, 1069) was added to certain wells. Five to ten min later, a MEA recording was performed again. The MEA plate was placed back to the incubator and incubated at 37°C, 5% CO_2_ for a specified number of hours (12, 24, 36, or 48 h). After that time, TTX was removed. Briefly, a first MEA recording was performed. All the spent medium was then aspirated. This washing step was repeated three times, in order to remove TTX completely. Then, 50% pre-conditioned untreated culture medium and 50% fresh medium were immediately added to the cells. At the last step, MEA recording was performed again. MEA plates were kept in the recording chamber for a longer measurement.

### MEA recordings and data analysis

All MEA recordings were performed using the 24-well MEA system (Multichannel Systems, MCS GmbH, Reutlingen, Germany). Recordings and data analysis procedures have been described previously in detail (Frega et al., 2019; Mossink et al., 2021). Briefly, spontaneous electrophysiological activity of hiPSC-derived neurons grown on MEAs was recorded for 10 min. Before recording, MEA plates were in exploring status for 10 min, in order to adapt in the recording chamber. After 10 min, recording started. During the recording, the temperature was maintained constantly at 37°C, and the evaporation and pH changes of the medium were prevented by inflating a constant, slow flow of humidified gas (5% CO_2_ and 95% O_2_) onto the MEA plates (with the lid on). The signal was sampled at 10 kHz, filtered with a high-pass filter (i.e., Butterworth, 100 Hz cutoff frequency), and the noise threshold was set at ± 4.5 standard deviations.

Data analysis was performed off-line by using the Multi-well Analyzer (i.e., software from the 24-well MEA system that allows the extraction of the spike trains) and a custom-made MATLAB (The Mathworks, Natrick, MA, USA) code that allows the extraction of parameters describing the network activity. Mean firing rate (MFR) was defined as the average of the spike frequency of all 12 channels across one well of the MEA plate. For the burst detection, the number of bursting channels (above threshold 0.4 burst/s and at least 5 spikes in one burst with a minimal inter-burst-interval of 100 ms) was determined. A network burst was called when at least half of the channels in one well presented a synchronous burst.

### Chemicals

TTX and 1-naphthyl acetyl spermine trihydrochloride (NASPM) were freshly prepared into concentrated stocks and stored frozen at −20°C. TTX was dissolved in distilled water (1 mM, Tocris 1069); NASPM was dissolved in distilled water (100 mM, Tocris 2766). For NASPM experiments on MEAs, immediately before adding NASPM to the cells, an aliquot of the concentrated stock was first diluted 1:100 at room temperature in Dulbecco’s phosphate-buffered saline (DPBS) and vortexed briefly. Then, the appropriate amount of working dilution (1:100, the final concentration of NASPM was 1 μM) was added directly to wells on the MEA after cells were refreshed with 50% pre-conditioned untreated culture medium and 50% fresh medium, and mixing was primarily through diffusion into the cell culture medium.

### Immunocytochemistry

Cells cultured on coverslips were fixed with 4% paraformaldehyde supplemented with 4% sucrose for 15 min at room temperature, followed by permeabilization with 0.2% Triton in PBS for 10 min at room temperature. Nonspecific binding sites were blocked by incubation in blocking buffer (PBS, 5% normal goat serum, 1% bovine serum albumin, 0.2% Triton) for 1 h at room temperature. Cells were incubated in a primary antibody solution wherein antibodies were diluted in blocking buffer overnight at 4°C. Secondary antibodies, conjugated to Alexa-fluorochromes, were diluted in blocking buffer and applied for 1 h at room temperature. Hoechst 33342 (Molecular Probes) was used to stain the nucleus before the cells were mounted using DAKO fluorescent mounting medium (DAKO). For the surface GluA2 staining and axon initial segment staining, neurons were fixed 48 h after TTX treatment at DIV 49. To ensure extracellular staining and prevent intracellular staining, permeabilization was not performed when using anti-GluA2. The primary antibodies that were used are: Mouse anti-GluA2 (1:500, Invitrogen 32-0300); Rabbit anti-MAP2 (1:1000, Abcam, #ab32454); Guinea pig anti-Synapsin 1/2 (1:1000, Synaptic Systems 106004); Mouse anti-Ankyrin G (1:200, Invitrogen 33-8800). The secondary antibodies that were used are: Goat-anti-mouse Alexa 488 (1:1000, Invitrogen A11029); Goat anti-Rabbit Alexa Fluor 488 (1:1000, Invitrogen A11034); Goat anti-Guinea Pig Alexa Fluor 568 (1:1000, Invitrogen A11075); Goat anti-Mouse Alexa Fluor 568 (1:1000, Invitrogen A11031); Goat anti-Rabbit Alexa Fluor 647 (1:1000, Invitrogen A21245). Cells were imaged at 63x magnification using the Zeiss Axio Imager Z1 equipped with ApoTome. All conditions within a batch were acquired with the same settings in order to compare signal intensities between different experimental conditions. Fluorescent images were analyzed using FIJI software. The number of GluA2 puncta was determined per individual cell via manual counting and divided by the dendritic length of the neuron. AIS start and end positions were obtained at the proximal and distal axonal positions.

### Whole cell patch clamp

Whole cell patch clamp was performed as previously described (Mossink et al., 2022). Vehicle treated- or 48 h TTX-exposed coverslips were placed in the recording chamber, continuously perfused with oxygenated (95% O_2_/ 5% CO_2_) artificial cerebrospinal fluid (ACSF) at 32°C containing (in mM) 124 NaCl, 1.25 NaH_2_PO_4_, 3 KCl, 26 NaHCO_3_, 11 Glucose, 2 CaCl_2_, and 1 MgCl_2_. Patch pipettes with filament (ID 0.86 mm, OD1.05 mm, resistance 6-8 MΩ) were pulled from borosilicate glass (Science Products GmbH, Hofheim, Germany) using a Narishige PC-10 micropipette puller (Narishige, London, UK). For all recordings, a potassium-based intracellular solution containing (in mM) 130 K-Gluconate, 5 KCl, 10 HEPES, 2.5 MgCl_2_, 4 Na2-ATP, 0.4 Na3-ATP, 10 Na-phosphocreatine, and 0.6 EGTA was used, with a pH of 7.2 and osmolality of 290 mOsmol/L. Miniature postsynaptic currents (mEPSCs) were measured in ACSF containing 1 µM TTX. Cells were visualized with an Olympus BX51WI upright microscope (Olympus Life Science, PA, USA), equipped with a DAGE-MTI IR-1000E (DAGE-MTI, IN, USA) camera. A Digidata 1440A digitizer and a Multiclamp 700B amplifier (Molecular Devices) were used for data acquisition. Data was acquired at 10 KHz (mEPSCs), and a lowpass 1 kHz filter was used during recording. Recordings were not corrected for liquid junction potential (±10 mV), and they were discarded if series resistance reached >25 MΩ or dropped below a 10:0 ratio of Rm to Rs. mEPSCs were analyzed using MiniAnalysis 6.0.2 (Synaptosoft Inc, GA, USA).

### RNA-Sequencing

Cells were treated with or without 1 µM TTX (Tocris, 1069) at DIV 49 and harvested at DIV 51. In all conditions, hiPSC-derived neurons were co-cultured with rat astrocytes. For RNA-Sequencing (RNA-seq), the prepared samples were sequenced on an Illumina NovaSeq 6000 S1 platform at an average depth of >50 million reads per sample using a read length of 2*100 base pairs. RNA from three biological replicates of vehicle-treated (Veh) and 48 h-TTX-treated (TTX) neurons (6 samples in total) were isolated using Quick-RNA Microprep kit (Zymo Research, R1051) according to the instructions of manufacturer. RNA quality was assessed using Agilent’s Tapestation system (RNA High Sensitivity ScreenTape and Reagents, 5067– 5579/80). RNA integrity number (RIN) values of all samples ranged between 8.3 and 8.7. Library preparation and paired-end RNA-sequencing were performed at CNAG-CRG, Centre for Genomic Regulation (CRG) (https://www.cnag.crg.eu/). Briefly, the RNA-Seq libraries were prepared with KAPA Stranded mRNA-Seq Illumina® Platforms Kit (Roche) starting from 500 ng of total RNA. The poly-A fraction enrichment was performed with oligo-dT magnetic beads, followed by the mRNA fragmentation. The strand specificity was achieved during the second strand synthesis performed in the presence of dUTP instead of dTTP. The blunt-ended double stranded cDNA was 3′adenylated, and Illumina platform-compatible adaptors with unique dual indexes and unique molecular identifiers (Integrated DNA Technologies) were ligated. The ligation product was enriched with 15 polymerase chain reaction (PCR) cycles. The libraries were sequenced on NovaSeq6000 (Illumina) in paired-end following the manufacturer’s protocol for dual indexing. Image analysis, base calling, and quality scoring of the run were processed using the manufacturer’s software Real Time Analysis, followed by generation of FASTQ sequence files.

### RNA-Seq data processing

RNA-seq reads were mapped against the hybrid human and rat reference genome (GRCh38+ Rnor6.0) with STAR/2.7.8a (Dobin et al., 2013) using ENCODE parameters. Gene quantification was performed with RSEM/1.3.0 (Li and Dewey, 2011) using the human gencode39 and rat ensembl104 annotations. Raw count matrices were loaded in R v4.2.1. For Ensemble IDs that mapped to same gene symbols, we only considered the IDs with highest expression per sample. Then, count data were normalized using size factors method of DESeq2 (Love et al., 2014), and lowly expressed genes (< 15 normalized counts in three or more samples) were filtered. Principal component analysis was performed on variance-stabilized transformed counts, and differential expression analysis was performed with DESeq2 using *lfcShrink* and “apeglm” method (Love et al., 2014). Genes were considered differentially expressed if TTX-treatment induced a Log2FC > 0.58, with a false discovery rate (FDR)-adjusted P value < 0.05. Hierarchical clustering of the samples based on differentially expressed genes (DEGs) was performed with *heatmap.2* function of gplots v3.1.3R package, using the Euclidean distance and “ward.D2” clustering method. Gene ontology (GO) enrichment analysis was performed on highly and lowly expressed genes after TTX-treatment independently, using the *gost* function of the R package gprofiler2 v0.2.1. Redundancy of enriched GO terms was accounted for with clustering analysis and aggregating terms with high semantic similarity, using the functions *calculateSimMatrix* and *reduceSimMatrix* with threshold=0.7 of the rrvgo v1.2.0 R package. We tested whether DEGs induced by TTX treatment were enriched for genes related to neurodevelopmental disorders (NDDs) and confident autism-related genes (Fu et al., 2022; Leblond et al., 2021), using one-tailed Fisher’s exact test and adjusting P values with the FDR method. All plots were generated with custom code based on ggplot2 v.3.4.0 functions in R.

### Quantification and Statistical analysis

The statistical analysis of the data was performed using GraphPad Prism 9 (GraphPad Software, Inc., CA, USA). We first determined whether data were normally distributed. Significance analysis was done for different experimental conditions by one-way ANOVA or two-way ANOVA with *post-hoc* Bonferroni correction when different cell lines, time lines, and drug-treated samples were included. Analysis was done using unpaired Student’s t tests when comparing two conditions at a single time point. Results with P values < 0.05 were considered as significantly different (*), P < 0.005 (**), P < 0.0005 (***), P < 0.0001 (****). Data is shown as mean ± standard error of the mean (SEM). All means, SEM, and test statistics are listed in Table S3.

## Data and code availability

The GEO accession number for the RNA-seq data in this paper is GSE225761.

## Author contributions

X.Y., B.F. and N.N.K. conceived the hypothesis and designed the experiments. N.N.K., B.F. and D.S. supervised the study. X.Y. and J.R.V.R. performed the experiments. X.Y., S.P. and J.R.V.R. analyzed the data. A.E.C., M.M., S.R. and E.J.H.v.H. assisted in data analysis. A.O. and C.S. assisted in experiments. X.Y., S.P. and J.R.V.R drafted the manuscript. All authors reviewed and edited the draft manuscript.

## Acknowledgements

The work was supported by funding from the European Community’s Horizon 2020 Programme (H2020/2014 – 2020) under grant agreement n° 728018 (Eat2beNICE) (to B.F.); ERA-NET NEURON-102 SYNSCHIZ grant (NWO) 013-17-003 4538 (to D.S.); China Scholarship Council 201906100038 (to X.Y.); ISCIII /MINECO (PT17/0009/0019) and FEDER (to A.E.C.); M.M. was supported by an internal grant from the Donders Centre for Medical Neurosciences of the Radboud University Medical Center.

