## Supplemental file for "A human *in vitro* neuronal model for studying homeostatic plasticity at the network level"

**Supplemental information**

Supplemental information includes the five supplemental figures, four supplemental tables, supplemental experimental procedures, and supplemental references.

**Supplemental items**

Supplemental Figure S1. Quantification of microelectrode array (MEA) parameters in Ctrl2 and Ctrl3 neurons. Related to Figure 1.
Supplemental Figure S2. Effect of 1-naphthyl acetyl spermine trihydrochloride (NASPM) treatment on the neuronal network activity. Related to Figure 2.
Supplemental Figure S3. Miniature excitatory postsynaptic current (mEPSC) activity in Ctrl 3 neurons. Related to Figure 3.
Supplemental Figure S4. Gene expression changes in tetrodotoxin (TTX)-treated neurons. Related to Figure 4.
Supplemental Figure S5. Astrocytes and neurons share some transcriptional changes in response to TTX-induced neuronal activity suppression. Related to Figure 5.

**Supplemental tables (Table S1-3 were added as separate excel files)**

Table S1: Tab 1, Results from differential expression analysis for human induced pluripotent stem cell (hiPSC)-derived neurons treated with TTX for 48 h, related to Figure 4. Tab 2, Results from gene ontology (GO) enrichment analysis for hiPSC-derived neurons treated with TTX for 48 h, related to Figure 4. Tab 3 and 4, Results from comparison between differentially expressed genes (DEGs) in hiPSC-derived neurons induced by TTX treatment and genes related to neurodevelopmental disorders (NDDs) as well as confident autism-related genes, related to Figure 4. Tab 5, Results from SYNGO analysis for human induced pluripotent stem cell (hiPSC)-derived neurons treated with TTX for 48 h, related to Figure 4.

Table S2: Tab 1, Results from differential expression analysis for rat astrocytes treated with TTX for 48 h, related to Figure 5. Tab 2, Results from GO enrichment analysis for rat astrocytes treated with TTX for 48 h, related to Figure 5. Tab 3 and 4, Results from comparison between DEGs in rat astrocytes induced by TTX treatment and genes related to NDDs as well as confident autism-related genes, related to Figure 5. Tab 5, Results from SYNGO analysis for rat astrocytes treated with TTX for 48 h, related to Figure 5.

Table S3: Statistics per figure, including all means, SEM and test statistics.

Table S4: List of quantitative polymerase chain reaction (q-PCR) primers used in this study (added in this manuscript), related to Figure 5.

**Supplemental experimental procedures**

hiPSC line origin and generation information
Gene expression analysis

### Supplemental items

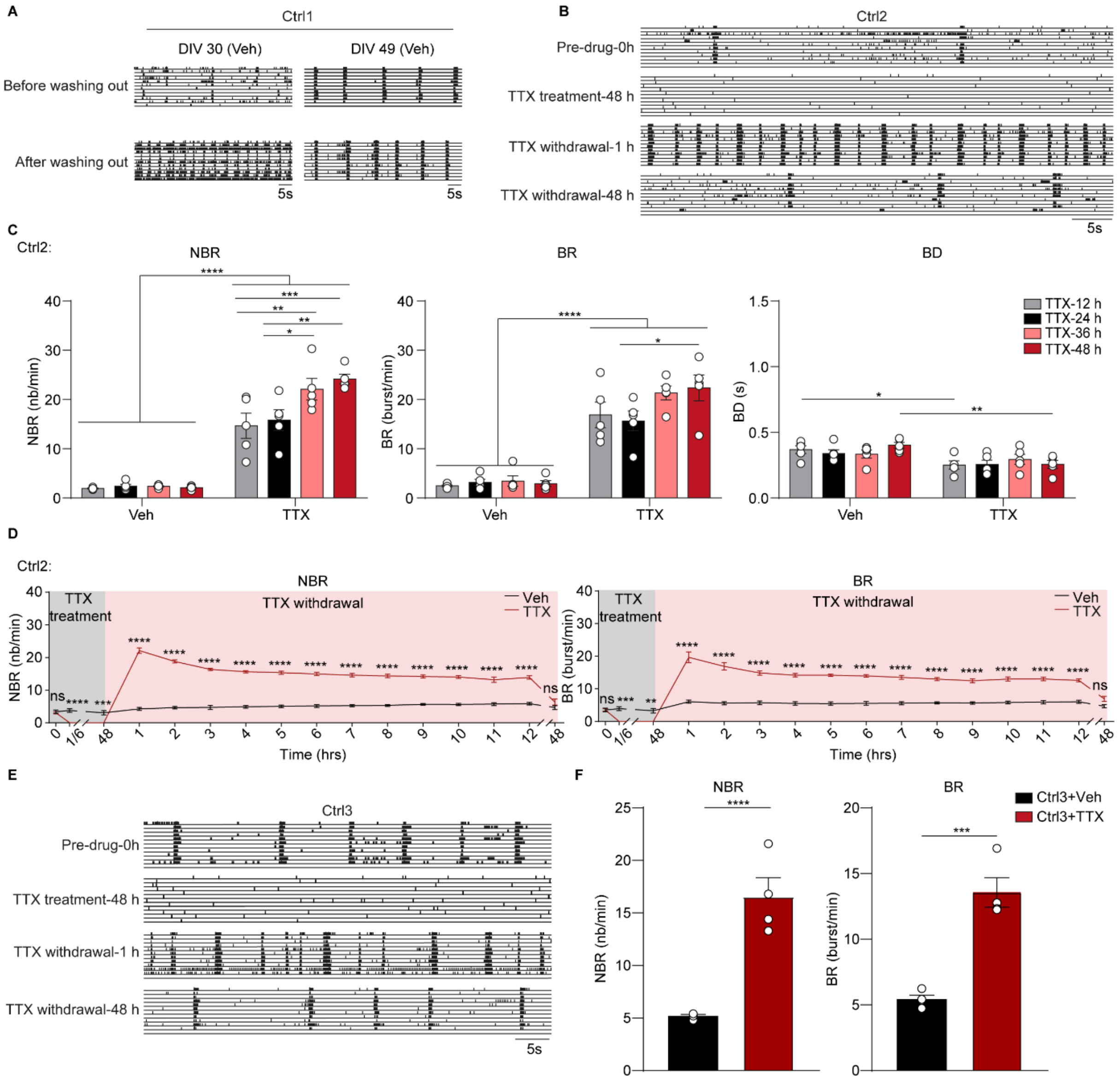


**Supplemental Figure S1. Quantification of microelectrode arrays (MEAs) parameters in Ctrl2 and Ctrl3 neurons. Related to Figure 1.**(A) Representative raster plots showing 1 min of spontaneous activity from hiPSC-derived neuronal networks (Ctrl1) before and after washing out procedure at days *in vitro* (DIV) 30 and DIV 49. (B) Representative raster plots showing 1 min of spontaneous activity from hiPSC-derived neuronal networks (Ctrl2) before and after TTX treatment, including before addition of TTX (Pre-drug-0 h), 48 h after addition of TTX (TTX treatment-48 h), 1 h after TTX withdrawal (TTX withdrawal-1 h), and 48 h after TTX withdrawal (TTX withdrawal-48 h). (C) Bar graphs showing the effect of 12 h, 24 h, 36 h, and 48 h TTX treatment on the mean network burst rate (NBR), mean burst rate (BR), and mean burst duration (BD) for Ctrl2 neuronal networks. All MEA parameters were measured 1 h after TTX withdrawal. n = 6 wells for all vehicle-treated (Veh) and TTX-treated (TTX) conditions. (D) Quantification of NBR and BR over time for vehicle-treated (Veh) and 48 h TTX-treated (TTX) neurons (Ctrl2). n = 8 wells for Veh, n = 10 wells for TTX. (E) Representative raster plots showing 1 min of spontaneous activity from hiPSC-derived neuronal networks (Ctrl3) before and after TTX treatment, including before addition of TTX (Pre-drug-0 h), 48 h after addition of TTX (TTX treatment-48 h), 1 h after TTX withdrawal (TTX withdrawal-1 h), and 48 h after TTX withdrawal (TTX withdrawal-48 h). (F) Bar graphs showing the quantification of NBR and BR for (E) at TTX withdrawal-1 h. n = 4 wells for Veh, n = 4 wells for TTX. Data represent means ± SEM. ns: not significant, $^{*}P < 0.05$, $^{**}P < 0.005$, $^{***}P < 0.0005$, $^{****}P < 0.0001$. For panel D, unpaired Student's T-test was performed between two groups. For panel C and D, two-way ANOVA test followed by a *post-hoc* Bonferroni correction was performed between conditions. For panel F and G, unpaired Student's T-test was performed between two groups. All means, SEM and test statistics are listed in Table S3.

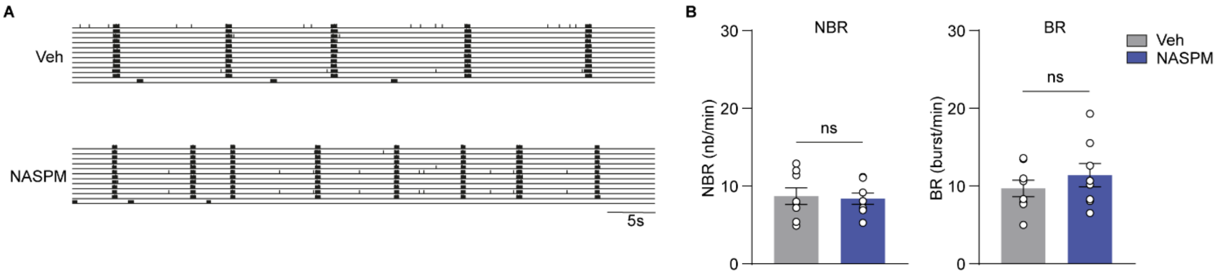


**Supplemental Figure S2. Effect of 1-naphthyl acetyl spermine trihydrochloride (NASPM) treatment on the neuronal network activity. Related to Figure 2.** (A) Representative raster plots showing 1 min of spontaneous activity from Ctrl1 neuronal networks grown on MEAs at DIV 49. Where indicated, the cells were vehicle-treated (Veh), or cells were treated with 10 µM NASPM before TTX application (NASPM). (B) Bar graphs showing the quantification of NBR and BR for (A). n = 8 for Veh, n = 8 for NASPM. Data represent means ± SEM. ns: not significant, unpaired Student's T-test was performed between two groups. All means, SEM and test statistics are listed in Table S3.

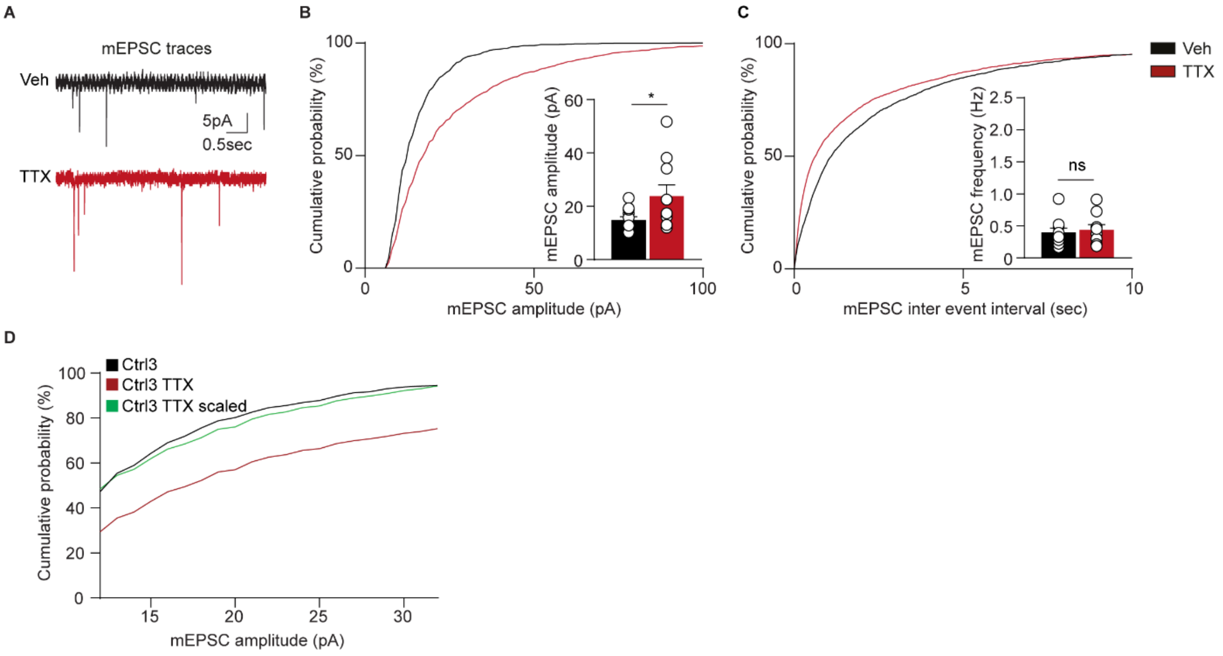


**Supplemental Figure S3: Miniature excitatory postsynaptic current (mEPSC) activity in Ctrl 3 neurons. Related to Figure 3.** (A) Representative mEPSC traces of vehicle-treated (Veh) and 48 h-TTX treated (TTX) neurons (Ctrl3). (B-D) Quantification of the amplitude (B) and frequency of mEPSCs (C) in Veh and TTX conditions. n = 11 recordings for Veh, n = 10 recordings for TTX (in total 3 batches for each condition). Rescaled cumulative mEPSC amplitude (Scaling = TTX values * 1.19) (D). Data represent means ± SEM. *$P < 0.05$, unpaired Student's T-test was performed between two groups. All means, SEM and test statistics are listed in Table S3.

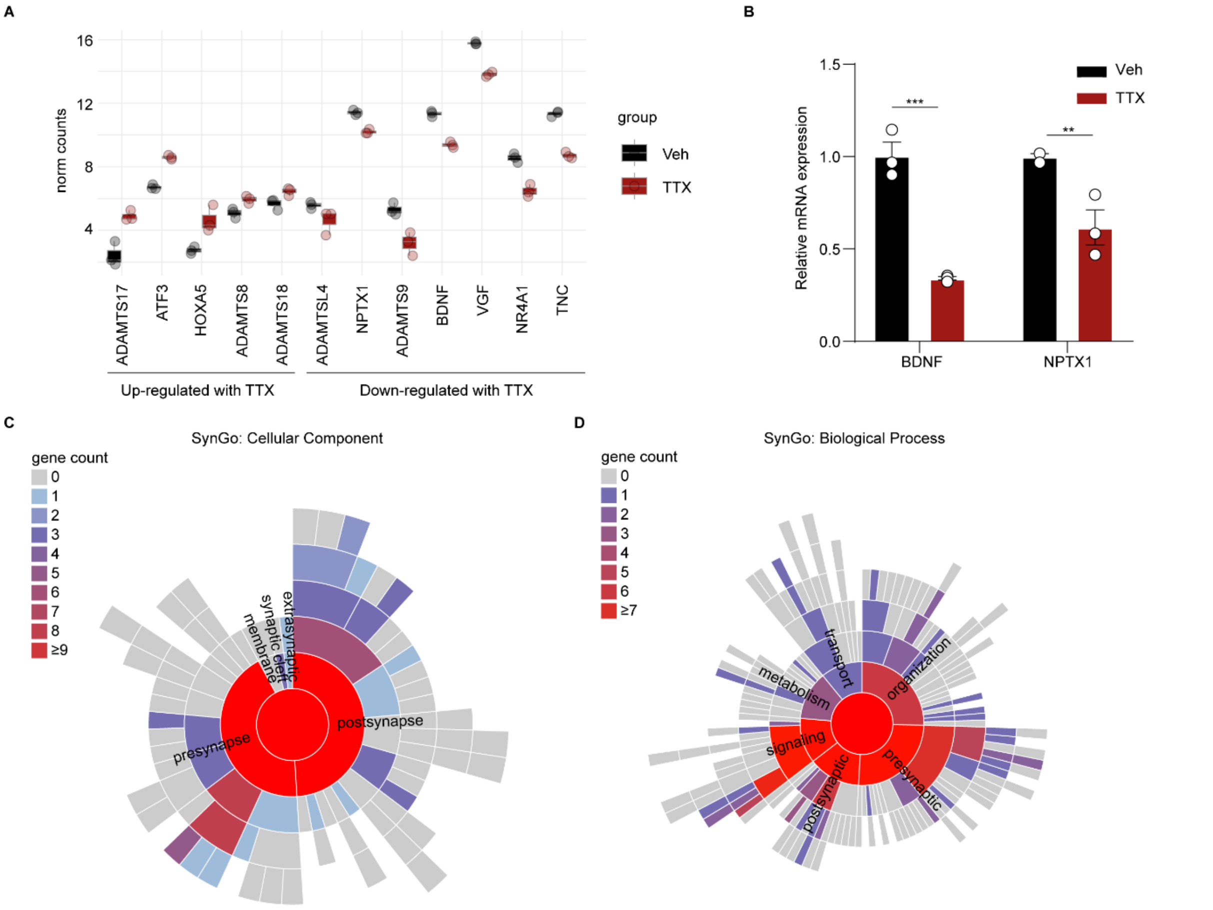


**Supplemental Figure S4. Gene expression changes in TTX-treated neurons. Related to Figure 4.** (A) Boxplot depicting the expression of selected DEGs (DESeq2 normalized counts), in both TTX- (red) and vehicle-treated neurons (black). (B) Bar graph showing relative messenger RNA (mRNA) expression of selected genes in TTX treated neurons (TTX-treated neurons = red, and vehicle-treated neurons = black). n = 3 replicates for each group. Data represent means ± SEM. $^{**}P < 0.005$, $^{***}P < 0.0005$, two-way ANOVA test followed by a *post-hoc* Bonferroni correction was performed between conditions. All means, SEM and test statistics are listed in Table S3. (C-D) Sunburst plots for synaptic GO analysis of DEGs in response to TTX treatment in hiPSC-derived neurons (Ctrl1, absolute $\log_2$ fold change ($Log_2FC$) > 0.58 and adjusted P value < 0.05), including Cellular Component (C) and Biological Process (D). 41 DEGs have a Cellular Component annotation, 30 DEGs for Biological Processes (a gene may have multiple annotations). 0 Cellular Component terms are significantly enriched at 1% FDR (testing terms with at least three matching input genes), 0 for Biological Processes. Enrichment analyses were performed using 1-sided Fisher exact tests (with “greater than” for the alternative hypothesis) and the False Discovery Rate (FDR) method was applied for multiple testing correction. Synaptic GO analysis was performed with the SynGo (Koopmans et al., 2019). Warmer colors represent the predominance of genes associated with the respective pathway.

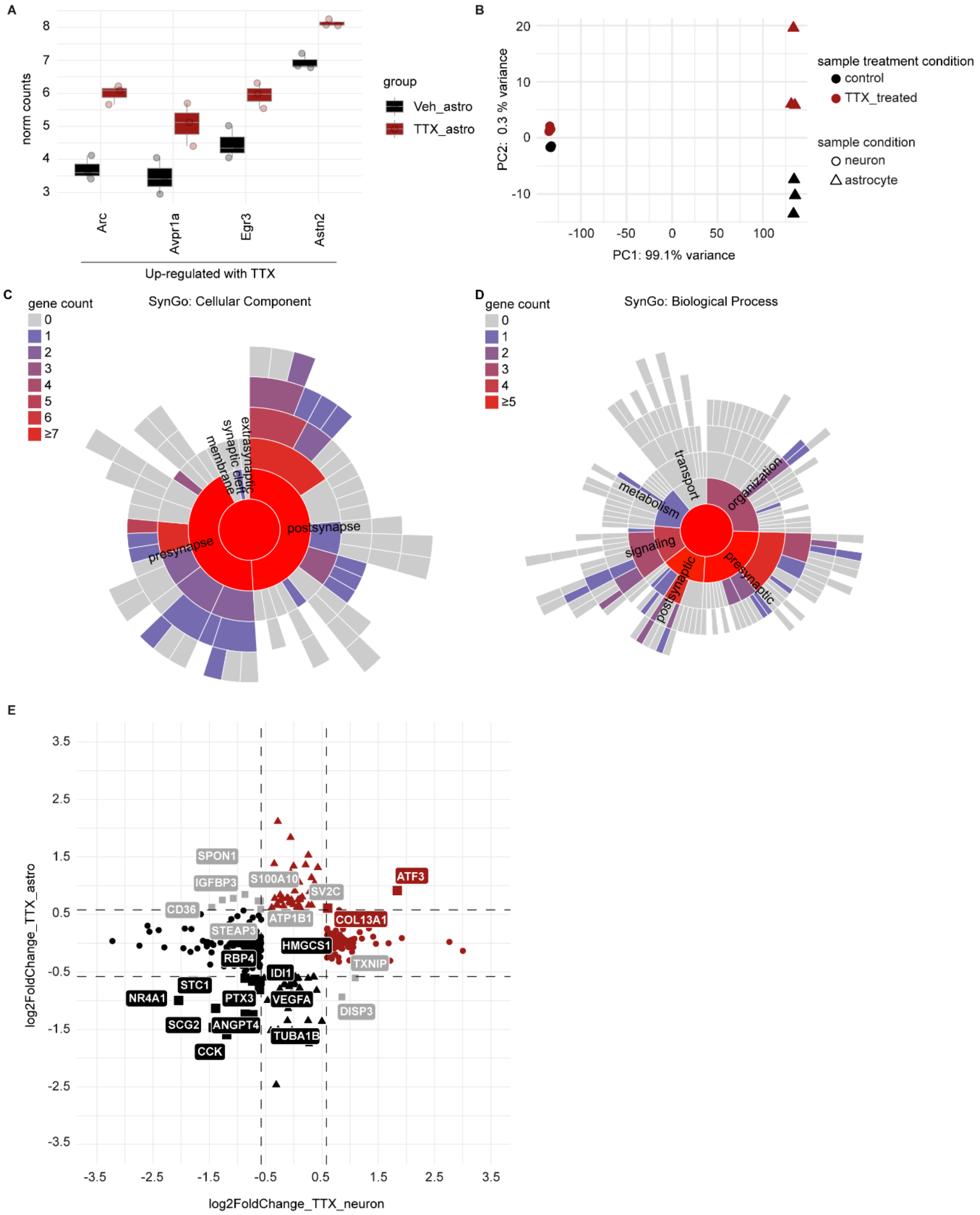


**Supplemental Figure S5. Astrocytes and neurons share some transcriptional changes in response to TTX-induced neuronal activity suppression. Related to Figure 5.** (A) Boxplot depicting the expression of selected DEGs (DESeq2 normalized counts), in both TTX- (red) and vehicle-exposed astrocytes (black). (B) Principal component analysis (PCA) of integrated RNA-sequencing counts from six samples of rat astrocytes and six samples of hiPSC-derived neurons. Shape of the points indicates the cell type (triangle = astrocyte, dot = neuron), and its colour depicts the received treatment (red = TTX and black = vehicle). (C-D) Sunburst plots for synaptic GO analysis of DEGs in response to TTX treatment in rat astrocytes co-cultured with hiPSC-derived neurons (Ctrl1, absolute $log_2$ fold change ($Log_2FC$) > 0.58 and adjusted P value < 0.05), including Cellular Component (C) and Biological Process (D). 20 DEGs have a Cellular Component annotation, 14 DEGs for Biological Processes (a gene may have multiple annotations). 2 Cellular Component terms are significantly enriched at 1% FDR (testing terms with at least three matching input genes), 0 for Biological Processes. Enrichment analyses were performed using 1-sided Fisher exact tests (with “greater than” for the alternative hypothesis) and the False Discovery Rate (FDR) method was applied for multiple testing correction. Synaptic GO analysis was performed with the SynGo (Koopmans *et al.*, 2019). Warmer colors represent the predominance of genes associated with the respective pathway. (E) Four-way volcano plot depicting expression changes per cell type in co-cultures of hiPSC-derived neurons and rat astrocytes treated with TTX (absolute Log2FC > 0.58 and adjusted p value < 0.05). Each data point represents a gene. Triangles depict DEGs only in astrocytes, circles indicate DEGs only in neurons, while squares depict DEGs in both astrocytes and neurons exposed to TTX. Color indicates the direction of change in gene expression activity in response to TTX treatment (red = up-regulated expression, black = down-regulated expression, gray = divergent direction in change in gene expression activity between the two cell types).

**Supplemental tables (Table S1-3 were added as separate excel files)**

Table S4. List of qPCR primers.

| Gene name | Direction | Primer sequence (5' to 3') |
|---|---|---|
| h*BDNF* | forward | GGATGAGGACCAGAAAGT |
| h*BDNF* | reverse | AGCAGAAAGAGAAGAGGAG |
| h*NPTX1* | forward | CACCGAGGAGAGGGTCAAGAT |
| h*NPTX1* | reverse | CAGGGCGGTTGTCTTTCTGA |
| h*PPIA* | forward | CATGTTTTCCTTGTTCCCTCC |
| h*PPIA* | reverse | CAACACTCTTAACTCAAACGAGGA |

### Supplemental experimental procedures

#### HiPSC line origin and generation information

Ctrl1 originated from fibroblasts of a 30-year old and healthy male doner (Mandegar et al., 2016; Miyaoka et al., 2014), and reprogrammed using episomal vector-based reprogramming of the Yamanaka transcription factors Oct4, c-Myc, Sox2, and Klf4 (Takahashi and Yamanaka, 2006), and was tested for genetic integrity using SNP assay (Frega et al., 2019). Ctrl2 was derived from fibroblasts of a 36-year-old and healthy female doner (Kondo et al., 2017; Okita et al., 2011), and reprogrammed using episomal vector-based reprogramming of the Yamanaka factors, showing no karyotypical malformations (Frega et al., 2019). Ctrl3 was previously derived from a healthy 51-year old male doner and reprogrammed using expression of Yamanaka factors by non-integrating Sendai virus, was tested for genomic integrity based on SNP array (Frega et al., 2019).

#### Gene expression analysis

Co-cultures of hiPSC-derived neurons and rat astrocytes were treated with or without 1 μM TTX (Tocris, 1069) at DIV 49 and harvested at DIV 51. RNA samples of vehicle- and TTX-treated conditions were isolated using Quick-RNA Microprep kit (Zymo Research, R1051) according to manufacturer's instructions.1 μg RNA was retro-transcribed into complementary strand of DNA (cDNA) by the iScript cDNA Synthesis Kit (Bio-Rad Laboratories, 1708891) according to the manufacturer's instructions. Quantitative real-time PCR (qRT-PCR) reactions were performed in QuantStudio Real-Time PCR systems by using GoTaq qPCR master mix 2x with SYBR Green (Promega, A6002) according to the manufacturer's protocol. We used human specific reference gene (*PPIA*) for our gene expression analyses. Used qRT-PCR primers were listed in Table S4. All samples were analyzed in triple in the same plate and placed in adjacent wells. Reverse transcriptase-negative controls and no template-controls were included in our procedures. The Ct value of every target gene was normalized against the Ct value of the reference genes [ΔCt = [Ct(target)-Ct(*PPIA*)]. The relative gene expression was calculated as $2^{-\Delta\Delta Ct}$ and used as fold change of gene expression when compared to corresponding control conditions [$2^{-\Delta\Delta Ct} = 2^{\Delta Ct(target)-\Delta Ct(control)}$].
